# Looking for pathways related to COVID-19 phenotypes: Confirmation of pathogenic mechanisms by SARS-CoV-2 - Host interactome

**DOI:** 10.1101/2020.11.03.366666

**Authors:** Francesco Messina, Emanuela Giombini, Chiara Montaldo, Ashish Arunkumar Sharma, Mauro Piacentini, Antonio Zoccoli, Rafick-Pierre Sekaly, Franco Locatelli, Alimuddin Zumla, Markus Maeurer, Maria R. Capobianchi, Francesco Nicola Lauria, Giuseppe Ippolito, COVID 19 INMI Network Medicine for IDs Study Group.

## Abstract

In the last months, many studies have clearly described several mechanisms of SARS-CoV-2 infection at cell and tissue level. Host conditions and comorbidities were identified as risk factors for severe and fatal disease courses, but the mechanisms of interaction between host and SARS-CoV-2 determining the grade of COVID- 19 severity, are still unknown.

We provide a network analysis on protein–protein interactions (PPI) between viral and host proteins to better identify host biological responses, induced by both whole proteome of SARS-CoV-2 and specific viral proteins. A host-virus interactome was inferred on published PPI, using an explorative algorithm (Random Walk with Restart) triggered by all the 28 proteins of SARS-CoV-2, or each single viral protein one-by-one. The functional analysis for all proteins, linked to many aspects of COVID-19 pathogenesis, allows to identify the subcellular districts, where SARS-CoV-2 proteins seem to be distributed, while in each interactome built around one single viral protein, a different response was described, underlining as ORF8 and ORF3a modulated cardiovascular diseases and pro-inflammatory pathways, respectively. Finally, an explorative network-based approach was applied to Bradykinin Storm, highlighting a possible direct action of ORF3a and NS7b to enhancing this condition.

This network-based model for SARS-CoV-2 infection could be a framework for pathogenic evaluation of specific clinical outcomes. We identified possible host responses induced by specific proteins of SARS-CoV-2, underlining the important role of specific viral accessory proteins in pathogenic phenotypes of severe COVID-19 patients.

## Introduction

Whilst COVID-19 predominantly affects the respiratory system, it is a multisystem disease, with a wide spectrum of clinical presentations from asymptomatic, mild and moderate, to severe, fulminant disease (1). Host conditions and comorbidities such as high age, obesity, diabetes, hypertension, organ damages, inflammation and coagulation dysfunctionality, were identified as risk factors for severe and fatal disease courses(2), but the mechanisms of interaction between host and SARS-CoV-2 that activate pathological pathways and determine severity, are still unknown.

In last months, many studies clearly described several mechanisms of SARS-CoV-2 infection at cell and tissue level. It was observed that the replication of SARS-CoV-2, as well as all +RNA viruses, occurs in the cytoplasm of the host cell, inducing a membrane rearrangement of rough endoplasmic reticulum (ER) membranes into double-membrane vesicles (3, 4). Moreover, NSP8 along with NSP7 and NSP12, which yield the RNA polymerase activity of NSP8, are assembled into the replicase–transcriptase complex, that begins to generate anti-sense (−) genomic RNA molecules, templates for positive-sense genome (+) and mRNA transcripts (5, 6). During virus entry, a well-known process is the cleavage of S protein by Furin on the cell membrane, which lead to split S protein into two subunits, S1 and S2, which the last can interact with ACE2 (7, 8). However, the SARS-CoV-2-host interaction is not restricted to local infection, but it triggers a systemic reaction, including the activation of the Bradykinin Storm, as described in many severe COVID-19 patients (9). Indeed, SARS-CoV-2 infection causes from one side a decrease of ACE level in the lung cells and, on the other side, an increase of ACE2 level, leading to increase Bradykinin level (9). Furthermore, microvascular injuries, due to systemic inflammatory response and endothelial dysfunction, were frequently found in severe COVID-19 patients (10). Increased circulating D-dimer concentrations, reflecting pulmonary vascular bed thrombosis with fibrinolysis, and elevated cardiac enzyme concentrations, linked to emergent ventricular stress induced by pulmonary hypertension, were associated to early features of severe pulmonary intravascular coagulopathy related to COVID-19 (11). Moreover, myocardial injuries were found to be linked to the risk of fatal outcome in COVID-19 patients (12–14). In 43% severe patients (21% of all COVID-19 patients) the myocardial injury was observed and a severe patient had a 4.74-fold increase in the risk of myocardial injury than non-severe patients (15). Although microvascular injuries and microthrombi formation frequently occurred in severe COVID-19 patients, the role of SARS-CoV-2 in this phenotype is not completely explained yet.

Recently, results of a meta-analysis study on interleukin-6 concentrations in patients with severe or critical COVID-19, pointed out that the systemic inflammatory profile of COVID-19 is distinct from that of non-COVID-19 ARDS, sepsis, and CAR T cell-induced cytokine release syndrome, and it might be involved in organ dysfunction in COVID-19, such as endovasculitis, direct viral injury and viral-induced immunosuppression (16).

Although many aspects of COVID-19 pathogenesis and mechanisms of SARS-CoV-2 have been investigated, only few papers describe the interactions among SARS-CoV-2 proteins by wet experiments. Physical associations in human cells, between SARS-CoV-2 proteins and human proteins, were found by affinity-purification mass spectrometry (AP-MS), identifying 332 high-confidence SARS-CoV-2-human protein-protein interactions (PPIs), highlighting an activation of innate immune pathways (17). Then, the influence of SARS-CoV-2 on transcriptome, proteome, ubiquitinome and phosphoproteome of a lung-derived human cell line, was described through a multi-omic approach. Such a multilevel representation highlighted autophagy mechanisms regulation by ORF3a and nsp6, the modulation of innate immunity by M, ORF3a and NS7b, and the Integrin-TGF-β-EGFR-RTK signalling perturbation by ORF8 of SARS-CoV-2 (18).

Virus–host interactome by computational approach has been applied to COVID-19 for drug repurposing (17), allowing the identification of new drug targets (19) and contributing to explain clinical manifestations (20, 21). The structural information on SARS-CoV-2 proteins and their interactions with human proteins and other viral proteins, allowed to better understand the mechanisms of SARS-CoV-2 infection, also comparing it with SARS-CoV (22).

One of the most interesting models of SARS-CoV-2 – host interactions was carried out on Spike-receptor interactions in other Human Coronaviruses (H-CoV). This network-based analysis of H-CoV - host interaction has provided a theoretic model for H-CoV infections applicable to SARS-CoV-2 pathogenesis, revealing biologically and clinically relevant molecular targets of three H-CoV infections (23).

The interactome based on PPI and gene expression data, have been applied also to uncover the molecular origins of phenotypes of other complex diseases (24, 25), in order to better define the mechanisms of COVID-19 pathogenesis.

The understanding of all mechanisms of SARS-CoV-2 infection also passes by overall visualization of biological reactions and pathways involved in COVID-19 and H-CoV infections. It can carry out on a disease map, such as COVID-19 Disease Maps, containing many diagrams about host molecular response during the infection (26–28).

In this context, defining the host response induced by specific viral proteins would be of great importance and can guide the identification of functional viral targets, helping to better define the pathologic phenotypes of the infection.

Here, we carry out a network analysis on PPI to better identify host biological response induced by SARS-CoV-2. Furthermore, the interactome analysis was applied to design network of SARS-CoV-2 - host proteins that could lead to a Bradykinin Storm. We used three different applications to identify interacting proteins: 200 proteins that interact closely with 28 SARS-CoV-2 proteins, 50 protein interactomes around each SARS-CoV-2 protein and 200 protein associated with KNG1, the pre-cursor of Bradykinin (BK).

In all three models, pathways associated with proteins were identified, including DNA damage/repair, TGF-β signalling, complement cascades (Figure 1).

**Figure 1.**
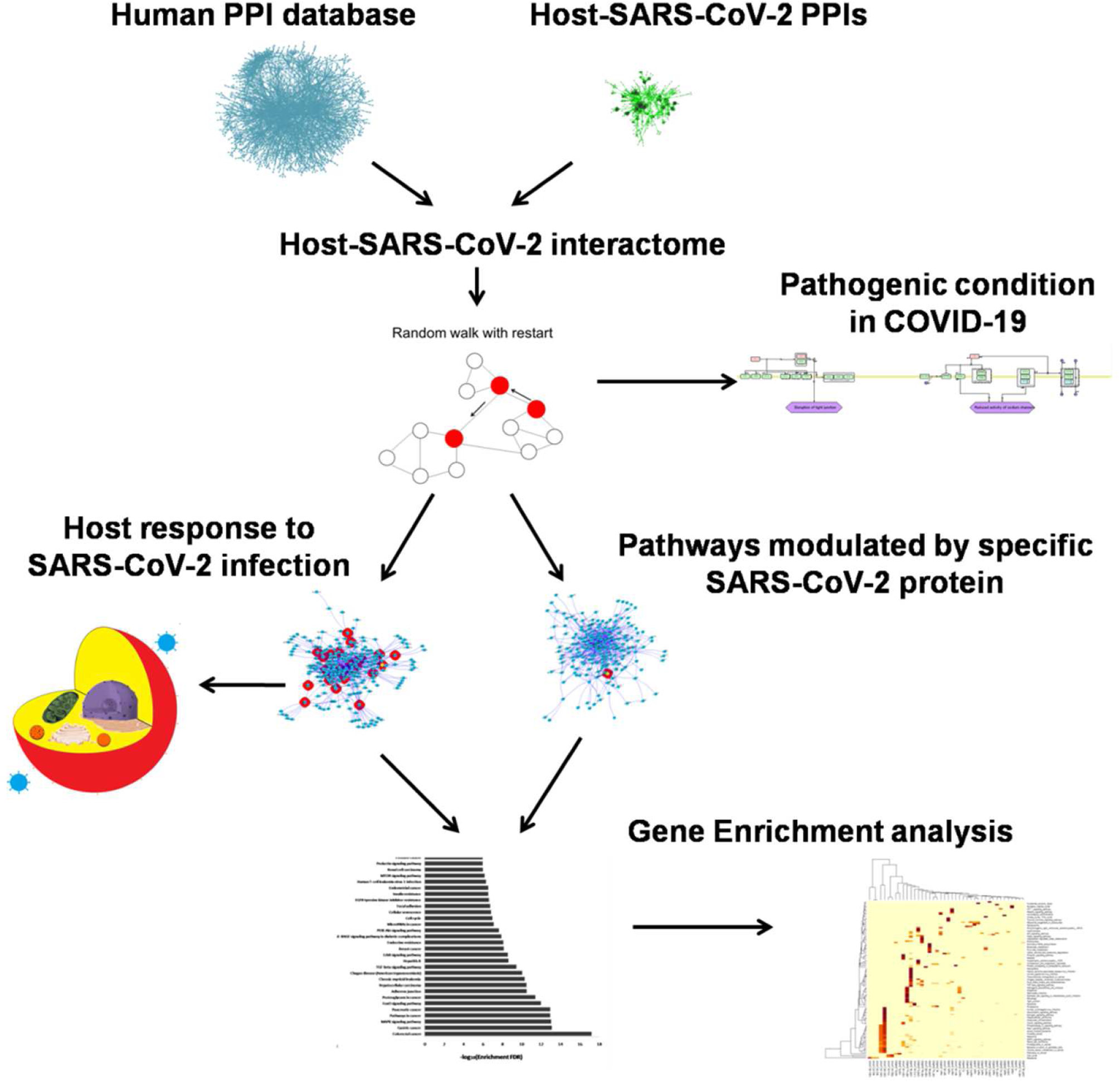
Workflow to describe SARS-CoV-2 host interactome analysis based on PPI data.

## Results

### Interactions with whole proteome of SARS-CoV-2

To define the massive effect of SARS-COV-2 infection on the host cell, an interactome between the entire set of SARS-CoV-2 proteins and host was carried out. Observing the colours’ distribution, it is possible to distinguish three different areas, corresponding to as many subcellular districts, where SARS-CoV-2 proteins seem to be distributed (Figure 2 a). In fact, N, nsp1, nsp4 and nsp15 are posed around nuclear proteins, while S, along with nsp5, nsp10, nsp12, nsp13, nsp14, nsp16, ORF1a and ORF9b, were among cytosolic and membrane proteins, at the bounder of the interactome. Protein M and many accessory and not-structural proteins (NS7b, nsp6, nsp7, ORF3a, ORF7a, ORF8, ORF10 and ORF14) are around mitochondrial and endoplasmic proteins.

**Figure 2.**
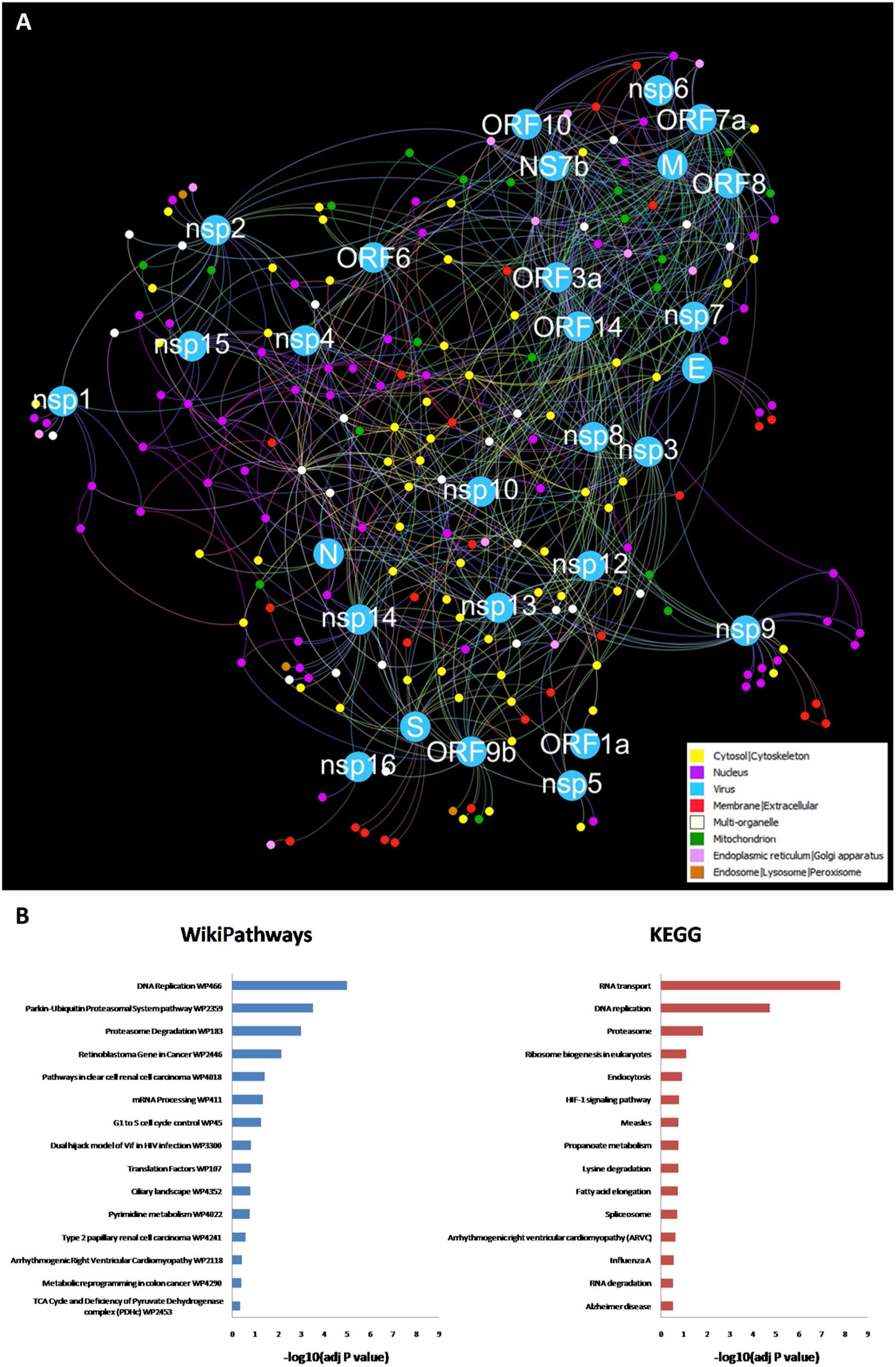
A) PPI interactome, based on human PPI and SARS-CoV-2-host interactions, with top 200 closest proteins identified by RWR, using together 28 proteins of SARS-CoV-2 as seeds for only one RWR run. Different colours of node and edges represent different locations in the cell, which are reported in the legend and in S-Table 1. This network was represented by Force Atlas algorithm. B) KEGG human pathway and WikiPathways Gene Enrichment analyses for 200 proteins identified by RWR algorithm using together 28 proteins of SARS-CoV-2.

We described the high values of betweenness centrality and degree for 9 host proteins: two proteins Multi-organelle Amyloid-beta precursor protein (APP) and dual-specificity protein phosphatase (PTEN); the membrane protein Sodium/potassium-transporting ATPase subunit alpha-1 (ATP1A1); two cytoplasmic proteins, 14-3-3 protein theta (YWHAQ) and Ubiquitin (UBC); two proteins of endoplasmic reticulum and Golgi apparatus, Sarcoplasmic/endoplasmic reticulum calcium ATPase 2 (ATP2A2) and Unconventional myosin-VI (MYO6); the mitochondrial ATP synthase subunit alpha (ATP5F1A); one nuclear receptor for export of RNAs, Exportin-1 (XPO1), suggesting their main role in this infection (S-Figure 1, S-Table 1). In order to further dissect the interactions with the entire proteome of SARS-CoV-2, enrichment analysis was carried out with WikiPathways and Kyoto Encyclopaedia of Genes and Genomes (KEGG) databases. WikiPathways gene enrichment analysis revealed biological pathways of DNA replication, ubiquitination and proteasome, with high significance (FDR <0.01%). In addition, KEGG pathway enrichment analysis revealed DNA and RNA replication pathways as the most significant pathways (FDR <0.01%), as well as signalling pathways and viral infection pathways (Figure 2 b).

### Host response induced by specific protein of SARS-CoV-2

To define the effect of specific viral proteins on host response, interactomes were built, choosing one specific viral protein as seed, imposing to find the 50 closest proteins. These analyses allowed to produce 28 restricted interactomes, which defined the strictest biological interactions associated to seed proteins both among human proteins and other viral proteins. In S-Figure 2 the interactomes for structural proteins were reported, while in S-Figures 3 and 4 the interactomes for accessory and non-structural proteins were plotted.

Most of the interactomes have few viral proteins into the reconstruct PPI network (i.e. M, nsp1, and ORF3a), while into other interactomes the viral seed protein shared own human target proteins with other viral proteins or interacts directly with other viral proteins, such as nsp3, nsp10, ORF6 and ORF10.

The lists of proteins for each reduced interactome were submitted to gene enrichment analysis on WikiPathways and KEGG pathway databases (S-table 2 and 3, respectively). For each single viral protein interactome, the top 5% with the smallest p values of gene enrichment analysis was selected, reporting for KEGG and WikiPathways in Figure 3 a and b, respectively.

**Figure 3.**
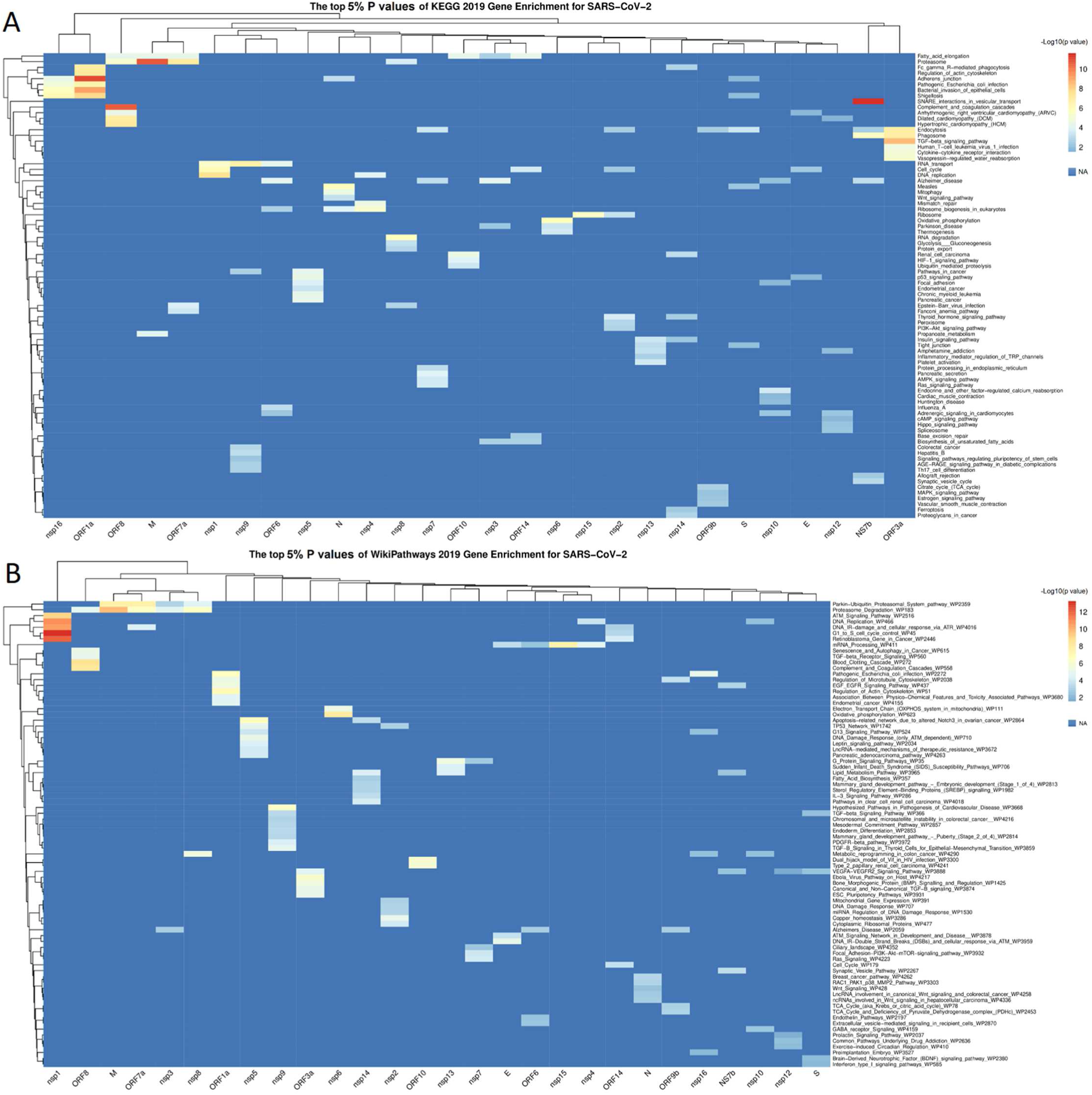
Heatmap reporting top 5% smallest P values of Gene Enrichment analysis on list of proteins, obtained by every interactome of single viral seed. The values were reported as -Log10 (p value). A) P values of Gene Enrichment based on KEGG 2019. B) P values of Gene Enrichment based on WikiPathways 2019.

For KEGG database the gene enrichment analysis on interactomes of NS7b, ORF1a, ORF3a and ORF8 showed pathway clusters highly significant and consistent with possible pathogenic mechanisms, such as the activation of the complement and of the coagulative cascade, (29) and the TGF-β-dominated immune response (30). In particular, the NS7b interactome revealed SNARE interactions in vesicular transport pathway, describing the mechanisms of intracellular vesicle trafficking and secretion, as the most significant pathway among all interactome (FDR< 0. 00001%). The ORF1a interactome revealed a cluster of pathways, composed by Adherents junction, Bacteria host infection mechanisms, Regulation of actin cytoskeleton and phagocytosis, all showing the direct involvement of this accessory proteins in viral entry in host cells (FDR < 0. 001%). The ORF8 interactome showed a pathway cluster involving complement and coagulation cascades and cardiovascular pathology (FDR < 0.3%). The ORF3a interactome revealed the TGF-β and Cytokine signalling, endocytosis and Vasopressin-regulation pathways (FDR < 0.01%). From pathway point of view, Proteasome pathway was involved in interactome of M protein as well as for ORF7a, ORF8 (FDR < 0.05%) (Figure 3a). For WikiPathways database the gene enrichment analysis showed results consistent with KEGG gene enrichment for ORF1a, ORF3a and ORF8, and added new pathway cluster for nsp1 (FDR< 0.00001%) with DNA replication and modification pathways (Figure 3b).

### Computational investigation on Bradykinin Storm in COVID-19

As the virus-host interactome has been efficient to describe the response to specific viral proteins, this method was applied to reconstruct the possible involvement of SARS-COV-2 proteins in triggering the Bradykinin Storm during the infection. In this case, Kininogen-1 (KNG1), precursor of bradykinin made by proteolysis cleavage, was considered as seed for RWR, imposing 200 closest proteins as limit to stop the algorithm. The resulting interactome showed NS7b, ORF3a, ORF8 and S as proximal to the seed protein, suggesting their role in the modulation of biological processes around Kininogen-1. Indeed, it would provide a direct influence of NS7b and ORF3a proteins to activation of BK. In fact, both NS7b and ORF3a interact with the same target, Endothelin Converter Enzyme (ECE1), and cell surface endopeptidase that converts big endothelin-1 to pressure peptide endothelin-1 (ET-1) and inactivates BK (Figure 4a). The enrichment analysis on WikiPathways and Kyoto Encyclopaedia of Genes and Genomes (KEGG) databases revealed biological pathways of Complement and coagulation cascades (FDR <0.000001%). Finally, to quantify the direct effect of S, ORF8, ORF3a, NS7b on Complement and coagulation cascades pathways, all proteins belonged to the most significantly enriched pathways, were highlighted on the interactome: Human Complement System WP2806 for WikiPathways (22 of 97 genes; adj P value 2.9E-21) and Complement and coagulation cascades for KEGG (24 of 79 genes; adj P value 5.96E-27). ORF8 showed the direct interactions with Fibrinogen alpha chain (FGA), described in both pathways, and Urokinase-type Plasminogen Activator (PLAU), Tissue-type Plasminogen Activator (PLAT) and Plasminogen activator inhibitor 1 (SERPINE1).

**Figure 4.**
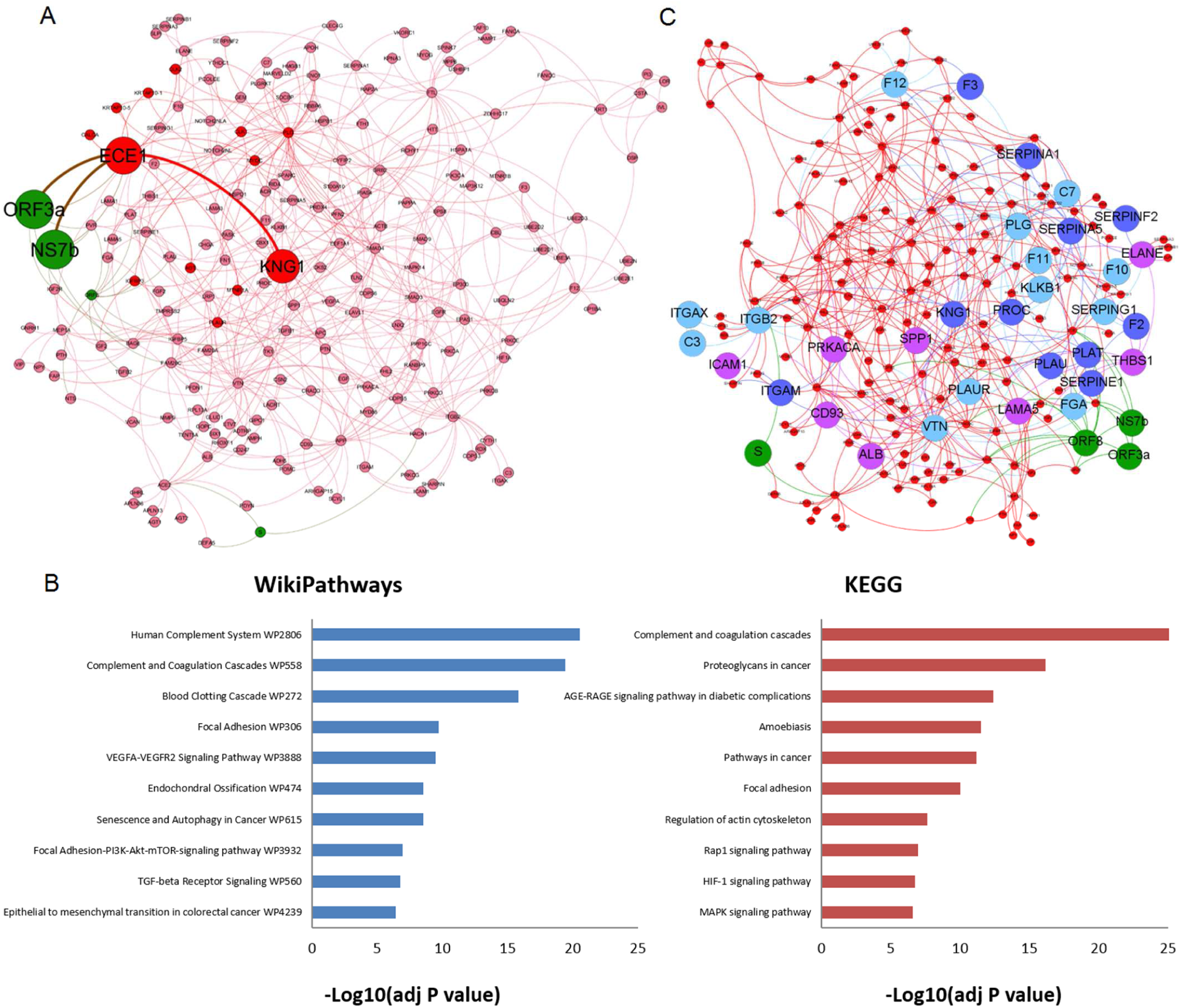
PPI interactome based on human PPI and SARS-CoV-2-host interactions, with top 200 closest proteins identified by RWR, using Kininogen-1 as seed. A) PPI Intercatome, where larger nodes and edges in red represent interactions via NS7b and ORF3a-ECE1-KNG1 (BK). The nodes in red are human proteins, while the nodes in green are virus proteins. B) KEGG human pathway and WikiPathways Gene Enrichment analyses for 200 proteins closest to KNG-1. C) PPI interactome, where proteins belonged to Human Complement System WP2806 (modes in purple) and Complement and coagulation cascades (nodes in blue) pathways were highlighted. The proteins belonged to both pathways were marked in light blue. The nodes in red are human proteins, while the nodes in green are virus proteins.

## Discussion

In this study, we built a functional interactome between SARS-CoV-2 and human proteome through RWR algorithm, to identify biological mechanisms and cell responses during SARS-CoV-2 infection and to propose a simulated model of infection contributing to a better understanding of COVID-19 pathogenesis. SARS-CoV-2 – host interactome allowed to mirror likely mechanisms of infections in different subcellular districts.

The high likelihood of our model to *in vivo* real mechanisms strengths the representative power of the interactome and the explorative algorithm to simulate biological processes.

Moreover, the proximity among specific viral proteins into the network, such as nsp7, nsp8, nsp9, nsp10, nsp12, nsp13, nsp14 involved in the viral RNA replication, can be a good way to investigate their possible function in viral infection. The closeness among S, nsp5, nsp16, ORF9b in the network is difficult to interpret, but S, nsp5 and ORF9b, along with N, had significant positive responses of IgG antibody in sera of COVID-19 patients (31). In SARS-CoV infected cell culture, the location of nsp2 in the cytoplasm and to some extent in the nucleus, as well as ORF3a, ORF7b, ORF6 and M in ER, seems to be consistent with their location among nuclear and cytoplasmatic, and ER proteins respectively (32).

The involvement of proteasome and ubiquitination pathways, along with RNA replication, represent the principal pathways activated for assembly and replication of SARS-CoV-2 (18). In fact, strong increases in RNA-modifying proteins were revealed in cell culture after infection with SARS-CoV-2 (33). The ubiquitin proteasome system deletes viral proteins to control the infection, but the virus can use them for its propagation (34).

The interactomes, built around a single protein of SARS-CoV-2, allowed to draw effects on cell, involving specific pathways. Vesicular transport mechanism by SNARE interactions, identified for NS7b, is used by pathogens to penetrate host cells through their membranes and in particular in SARS-CoV (35, 36).

The pathways showed in ORF3a and ORF8 interactomes suggested a possible modulation of coagulation cascade and cardiovascular pathology in COVID-19 and the involvement in Cytokine storm and water homeostasis in endothelial tissue, respectively.

Binding of ORF3a and ORF8 to TGF-β-associated factors (TGFB1, TGFB2, LTBP1, TGFBR2, FURIN, BAMBI) supports a strong involvement to begging pro-inflammation state, while the direct interactions between ORF8 and components of Coagulation pathway, such as FGB, FGA, C5, PLAU, PROS1, SERPINE1, PLAT and CLU (17, 18) suggest a modulation of coagulation cascade by SARS-CoV-2.

Although the biological function of the ORF8 protein of SARS-CoV-2 remains unclear, the role of ORF8 in severe COVID-19 outcome might be supported by SARS-CoV-2 variant with a 382-nucleotide deletion (Δ382) found in Singapore in January-February 2020 linked to mild forms of COVID-19. This deletion truncates open reading frame 7b and locks ORF8 transcription and would be associated to less cytokine releasing during the acute phase of infection (37, 38).

The systemic inflammatory response and the relied endothelial dysfunction might be responsible for the microvascular injuries resulting in the formation of microthrombi, as well as a featuring microvascular thrombosis and hemorrhage pathology, viewed in lung of COVID-19 patients (11). The main pathways described in ORF3a and ORF8 interactomes, are consistent with the microvascular injuries and the development of cardiovascular complications i. e. heart failure, myocarditis, pericarditis, vasculitis and cardiac arrhythmias observed in COVID-19 patients(39). Such featuring symptoms, suggest specific mechanisms of host-virus interactions, and reinforce the role of ORF3a and ORF8 in the COVID-19 pathogenesis.

To study the pathogenesis of COVID-19 in systemic context, Bradykinin Storm was investigated by the interactome approach. We showed NS7b and ORF3a interacting with ECE1, which can inactivate BK (40). Such observation suggests specific mechanisms of host-virus interactions, occurring in severe COVID-19 patients and reinforce the role of ORF3a to enhance the bradykinin dysregulation.

These host-virus interactions could enhance a better viral fitness during the infection, as suggested by the interaction already observed with SARS-CoV between NS7b in ECE1(18).

This interaction system might suggest a hypothesis that can explain the possible enhancing effect on Bradykinin Storm in COVID-19: the locking of ECE1 activity due to NS7b and especially ORF3a interaction, could reduce the activation of the vasoconstrictor endotelin-1, and amplify the vasodilatation effect of BK. The ECE1 gene is much more expressed in all the tissues than ACE2 and ACE, especially in the lungs, where ECE1 is 128 fold and 3.8 fold expressed compared to ACE2 and ACE, respectively (41).

Similar to the angiotensin peptides, BK is produced from an inactive pre-protein Kininogen-1 through the activation by the serine protease kallikrein (42). Furthermore, the excess of BK can lead to vasodilatation, hypotension and hypokalaemia (9, 43), which is associated with arrhythmia (44, 45). All these clinical conditions have been widely reported in COVID-19 patients (14, 46, 47). It is also notable that many of the other symptoms reported for COVID-19 (myalgia, fatigue, nausea, vomiting, diarrhoea, anorexia, headaches, decreased cognitive function) are remarkably similar to other hyper-BK-conditions that lead to vascular hyperpermeabilization (48). Moreover, the activation of BK is strongly linked to coagulation: active F12 activates plasma kallikrein (PK), that liberates bradykinin by cleavage (49). The direct interactions between ORF8 and two plasminogen activators, PLAU and PLAT, and Plasminogen activator inhibitor 1, SERPINE1, bulk proteins of Plasminogen Activator System and involved in lung injuries and neuroinflammation (31, 32), would suggest a modulating effect of ORF8 on Bradykinin Storm.

Moreover, inflammatory conditions highly induce an up-regulation of Bradykinin receptor B1 (BKB1R) (50). The results of our network-based model for pathogenesis of SARS-CoV-2 mirror likely clinical manifestations in severe COVID-19 patients, noting that analysis of published data about COVID-19 allows to synthesise and describe complex aspect of pathogenesis, as reported for inflammatory cytokines storm in COVID-19-induced respiratory failure (16).

### Limitations

There are many experimental platforms for deriving such physical interactions, such as Affinity purification mass-spectrometry (AP-MS) and yeast-two-hybrid (Y2H), which enable the accurate identification of interactions with a relatively long time.

The scenario reported in this study refers to few experimental data available on public databases and could be different respect to real phenotypes of COVID-19 patients.

The pathways’ analysis did not consider tissue and cell type diversity. Finally, the low threshold established for the number of nodes found by RWR (200) limited the reconstruction of the entire pathways. However, this was a software-imposed threshold. Although such network-based approach showed great potential in identifying mechanisms not yet observed, experimental tests will be necessary to confirm what we have described.

## Conclusion

We developed a network-based model for SARS-CoV-2 infection, which could be a framework for pathogenic evaluation of specific clinical outcomes.

Here, the PPI interactomes were used to identify mechanistic processes of the viral molecular machinery during SARS-CoV-2 infection, for viral survival and replication within the host. With this knowledge, proteins interactions crucial for pathogenesis could be discerned. We identified different host response induced by specific proteins of SARS-CoV-2, underlining the important role of ORF3a and ORF8 in phenotypes of severe COVID-19 patients. The interactome approach applied to identification of biological reactions around Kininogen-1, allowed to view NS7b and ORF3a interactions with ECE1, which might have a role to enhance Bradykinin storm. This network-based model of SARS-CoV-2 - host interaction could guide to develop novel treatments against specific viral proteins, such as monoclonal antibodies.

## Methods and Materials

The virus–host interactome was made by merging SARS-CoV-2 –host protein-protein interaction (PPI) data from Intact (51), with data from human PPI databases, such as BioGrid, InnateDB-All, IMEx, IntAct, MatrixDB, MBInfo, MINT, Reactome, Reactome-FIs, UniProt, VirHostNet, BioData, obtained by R packages PSICQUIC and biomaRt (52, 53). For SARS-CoV-2 - host interaction experimentally obtained, 1407 protein interactors and 2305 interactions were download from IntAct at 12 September 2020. 28 SARS-CoV-2 proteins are used for all the analyses: E, M, N, S, nsp1, nsp2, nsp3, nsp4, nsp5, nsp6, nsp7, nsp8, nsp9, nsp10, nsp12, nsp13, nsp14, nsp15, nsp16, ORF1a, ORF3a, ORF6, ORF7a, NS7b, ORF8, ORF9b, ORF10, ORF14.

To obtain functional information about PPI ex vitro, to simulate biological reactions in infected cells and to explore cell response against viral infection, we employed a Random walk with restart (RWR) algorithm, a state-of-the-art guilt-by-association approach(54). This algorithm allows to establish a proximity network from a given protein (seed), to study its functions, based on the premise that nodes related to similar functions tend to lie close to each other in the network. For this study, two types of interactome were performed: the whole SARS-CoV-2 -host interactome and an interactome per single SARS-COV-2 protein.

In the first imputation, every protein of SARS-CoV-2 proteome was used as seeds, with the limit of 200 closest host’s proteins to every SAR-CoV-2 protein, every protein of SARS-CoV-2 proteome was chosen as seeds for Random walk with restart, imposing the limit of 200 closest host proteins to every SARS-CoV-2 proteins. This analysis allowed to design SARS-COV-2-host interactome, where the different colours correspond to the localization of the proteins within the cell.

For the second kind of interactome, one SARS-CoV-2 per time was used as seed, lowering to closest 50 proteins in order to define the induced biological response as better as possible.

For each node, a score was computed as a measure of proximity to the seed protein (23). In total, a large PPI interaction database was assembled, including 13334 nodes and 73584 interactions. Graphical representations of networks were performed by GEPHI 0. 9. 2 (55). To identify hub protein in the SARS-CoV-2 - host interactome, the values of betweenness centrality and degree were plotted. Betweenness centrality score measure how a specific node is in-between other nodes and then can be considered a hub, while the degree of node corresponds to number of connections.

Pathways of proteins involved in host response were tested by gene enrichment analysis on Kyoto Encyclopaedia of Genes and Genomes (KEGG) human pathways and WikiPathways databases (56). To allow gathering of results for every running, the R package enrichR was used, an R interface to web-based tool ‘Enrichr’ for analysing gene sets (57). The Enrichr analysis was performed using these statistical parameters: p-value (Fisher exact test), q-value (adjusted p-value for false discovery rate, FDR). Results for KEGG and WikiPathways were considered significant with a revised p-value < 0. 05. To infer pathways involved in single viral interactome, gene enrichment analyses for each viral interactome were collected along with p-values, as reported in enrichR package output. P-values of every single enrichment analyses were transformed by the function x = − log10 (p-value) and the 5% of x values were plotted on heatmap by R package pheatmap (58).

## Acknowledgments

We gratefully acknowledge: Collaborators Members of INMI COVID-19 study group; COVID 19 INMI Network Medicine for IDs Study Group: Abbate Isabella, Agrati Chiara, Al Moghazi Samir, Ascoli Bartoli Tommaso, Bartolini Barbara, Capobianchi Maria Rosaria, Capone Alessandro, Goletti Delia, Rozera Gabriella, Nisii Carla, Gagliardini Roberta, Ciccosanti Fabiola, Fimia Gian Maria, Nicastri Emanuele, Giombini Emanuela, Lanini Simone, D’Abramo Alessandra, Rinonapoli Gabriele, Girardi Enrico, Montaldo Chiara, Marconi Raffaella, Addis Antonio, Maron Bradley, Bianconi Ginestra, De Meulder Bertrand, Kennedy Jason, Khader Shabaana Abdul, Luca Francesca, Maeurer Markus, Piacentini Mauro, Merler Stefano, Pantaleo Giuseppe, Rafick-Pierre Sekaly, Sanna Serena, Segata Nicola, Zumla Alimuddin, Messina Francesco, Vairo Francesco, Lauria Francesco Nicola, Ippolito Giuseppe.

## Abbreviations

PPI: Protein-Protein Interactions
SARS-CoV: Severe Acute Respiratory Syndrome Coronavirus
SARS-CoV-2: Severe Acute Respiratory Syndrome Coronavirus 2
H-CoV: Human Coronavirus
COVID-19: Coronavirus Disease 19
ORF: Open Reading Frame
RWR: Random walk with restart
FDR: False Discovery Rate
ER: Endoplasmic reticulum
Nsp: non-structural protein

## Authors’ contributions

Conceptualization: F. M., F. N. L. Data curation: F. M., E. G., Formal analysis: F. M. Funding acquisition: M. R. C., G. I. Investigation: F. M., F. N. L. Methodology: F. M. Resources: M. R. C., G. I. Software: F. M., Supervision: F. N. L. Validation: F. M. Visualization: F. M. E. G. Writing ± original draft: F. M., E. G., F. N. L. Writing ± review & editing: F. M., E. G., M. R. C., F. N. L, C. M. All authors reviewed and approved the manuscript.

## Author declarations

All authors have an interest in infectious diseases epidemics and public health. Authors declare no conflicts of interest.

## Consent for publication

Not applicable.

## Ethics approval and consent to participate

Not applicable.

## Funding

National Institute for Infectious Diseases Lazzaro Spallanzani–IRCCS received financial support funded by Italian Ministry of Health, grants: Ricerca Corrente program 1-Emerging and re-emerging infections; It-IDRIN, CCR-2017-23669075; and by projects COVID-2020-12371675 and COVID‐2020‐12371817.

This work was supported also by Findus Italia, part of the Nomad Foods.

Giuseppe Ippolito and Alimuddin Zumla are co-Principal Investigators of the Pan-African Network on Emerging and Re-Emerging Infections (PANDORA-ID-NET – https://www.pandora-id.net/) funded by thefunded by the European and Developing Countries Clinical Trials Partnership the EU Horizon 2020 Framework Programme. Sir Zumla is in receipt of a National Institutes of Health Research senior investigator award and is a Mahathir Science Award Laurete.

S-Table 1: Table with human and viral proteins identified by RWR in all SARS-CoV-2 proteins interactome are reported, along with statistical parameters and biological information.

S-Table 2: Table of gene enrichment analysis obtained by KEGG 2019 database. Results were considered significant with a p-value < 0. 05.

S-Table 3: Table of gene enrichment analysis obtained by WikiPathways 2019 database. Results were considered significant with a p-value < 0. 05.

S-Table 4: All Interactions selected by RWR in interactome for single seed viral protein. Detailed information about interactors, interaction and database were reported.

S-Figure 1. Plot betweenness centrality vs. degree of 199 human proteins, retained in all proteins of SARS-CoV-2 - host interactome. Limits were fixed to top 5% of Betweenness Centrality values and Degree, 0. 020554 and 11 respectively.

S-Figure 2. Interactomes based on human PPI and SARS-CoV-2-host interactions, with top 50 closest proteins identified by RWR, using structural proteins (M, S, N, E) as seed. The nodes in red are human proteins, while the nodes in green are virus proteins.

S-Figure 3. Interactomes based on human PPI and SARS-CoV-2-host interactions, with top 50 closest proteins identified by RWR, using accessory proteins (ORF1a, ORF3a, ORF6, ORF7a, NS7b, ORF8, ORF9b, ORF10, ORF14) as seed. The nodes in red are human proteins, while the nodes in green are virus proteins.

S-Figure 4. Interactomes based on human PPI and SARS-CoV-2-host interactions, with top 50 closest proteins identified by RWR, using accessory proteins (nsp1, nsp2, nsp3, nsp4, nsp5, nsp6, nsp7, nsp8, nsp9, nsp10, nsp12, nsp13, nsp14, nsp15, nsp16) as seed. The nodes in red are human proteins, while the nodes in green are virus proteins.

## Notes

### Competing Interest Statement

The authors have declared no competing interest.

